# Perfusion estimation using MRI-based measurements and a porous media flow model

**DOI:** 10.1101/2023.04.25.538213

**Authors:** Rolf J. Lorentzen, Geir Nævdal, Ove Sævareid, Erlend Hodneland, Erik Hanson, Antonella Z. Munthe-Kaas

**Affiliations:** NORCE Norwegian Research Centre AS, Bergen, Norway; Mohn Medical Imaging and Visualization Centre, Department of Radiology, Haukeland Universitetssykehus, Bergen, Norway; Department of Mathematics, University of Bergen, Bergen, Norway

## Abstract

The measurement of perfusion and filtration of blood in biological tissue give rise to important clinical parameters used in diagnosis, follow-up, and therapy. In this paper we address techniques for perfusion analysis using processed contrast agent concentration data from dynamic MRI acquisitions. New methodology for analysis is evaluated and verified using synthetic data generated on a tissue geometry.

**Author summary:** Accurate knowledge of tissue perfusion is crucial for proper diagnostics and treatment of several medical disorders. Traditional methods based on medical imaging are fast, but usually lack precision and robustness. In this paper, we address methodology to develop better diagnosis and treatment strategies for malignant tumors and stroke where blood perfusion may be altered. In our work, mathematical models for estimating perfusion are calibrated using magnetic resonance imaging (MRI) data, and more accurate representations of tissue parameters are provided. This methodology is a step towards minimal invasive and individually tailored diagnosis and treatment. We demonstrate the methodology with a twin-experiment using models of different complexity for generating data and estimating the tissue parameters. Both models are based on a mathematical description of how fluids flows in a porous medium, where the data-generating model uses higher resolution and a network representation of blood vessels than the estimating model. The calibration of unknown tissue parameters is done using a statistical framework, and the choice of methodology is motivated by applications from sub-surface reservoir characterization.

## Introduction

Accurate estimates for blood perfusion are of high importance in clinical applications. Perfusion imaging is used in treatment planning for stroke [1], but also for clinical applications related to staging of cancer [10].

Medical image acquisition techniques like computerized tomography (CT), magnetic resonance imaging (MRI), or positron emission tomography (PET), can all be applied in a dynamic setting where the evolving distribution of an injected contrast agent is gathered together as a temporal sequence of images. Quantitative tissue characterization (e.g. blood perfusion) from such data is traditionally performed locally by applying tracer-kinetic methodology [21] to a single region of interest (ROI) or a voxel at a time. While these approaches have advantages in terms of computational cost, they are generally considered to have certain systematic weaknesses, and the resulting estimates will typically lack robustness in terms of accuracy and reproducibility, see e.g. [8, 21]. One major problem is that the traditional methods are built around systems with assumed well defined boundaries and known contrast inflow distribution given as an arterial input function (AIF). This is a valid assumption if whole organs are considered, but does not necessarily hold when smaller regions of interest are investigated. It can be difficult to isolate the relevant arterial influx concentration, and the observed contrast agent concentration for the voxel will typically reflect an aggregation of flow in transit in the arterial and/or venous system as well as flow relevant to the local tissue perfusion or extravascular leakage. Local temporal deconvolution also neglects the potential useful spatial structure of the images.

In this work we will utilize the large potential to transfer methodology developed for the petroleum industry, to the medical sciences. In particular, advanced fluid models and numerical simulators developed for subsurface flow, can be used to predict the blood flow in organs. Models for flow in reservoir rocks have a large number of poorly known parameters, such as the spatial porosity and permeability (the rocks ability to transmit fluids). This is also the case when modeling blood flow in organs, as exact tissue properties are impossible to know, and are also subject to large variations between different individuals. Ensemble-based data assimilation is a popular approach to improve models for fluid flow, based on available measurements (see e.g. [3]). Iterative ensemble smoothers (see e.g. [18, 22]) have become the preferred approach for assisted history matching of petroleum reservoir properties. Smoothers assimilate all data at one single update step, and complicated and time consuming restarts of the simulator are avoided. However, a problem with ensemble based methods is that with a limited ensemble size, spurious correlations between observations and model parameters are inevitable. This problem increases with the size of the parameter space and the number of observations, and leads to unrealistic updates of model parameters, and underestimation of the posterior error covariance, see [7]. As a remedy for this, localization may be introduced in order to limit updates of parameters that we know are unlikely to influence a given observation. In our study we suggest to apply a correlation based localization, [17], where a threshold for the spurious correlations between the model parameters and the concentration measurements are estimated using the ensemble at the current iteration.

In this paper we investigate the potential of using fluid flow models in combination with an ensemble smoother for improved estimation of blood perfusion from dynamic MRI. Unknown parameters in a porous media flow model (based on Darcy’s equation) are updated, using a time sequence of spatially distributed contrast concentration data. Synthetic data are generated using a hybrid-scale model where flow in the largest vessels are simulated using Hagen-Poiseuille equation for laminar incompressible flow, and the tissue flow is modeled using Darcy’s equation. This hybrid-scale model is more complex compared to the pure porous media model, and the goal is to reconstruct the blood flow and perfusion despite the gap in complexity between the two models. The synthetic data have known, ground truth, perfusion maps. The methodology is tested on a domain representing a stretched-out frog tongue, with known vascular structure. A comparison is made with the classical maximum slope method and deconvolution based on singular value decomposition.

## Methods

We introduce two different modeling approaches. One to be used in parameter estimation and a more complex and computationally demanding to be used to generate synthetic data with known ground truth. For the purpose of data assimilation we consider a slightly modified version of the traditional dual-porosity, dual permeability model (see e.g., [2]). This partitioning of the geometry into multiple compartments allow a explicit characterization of the local tissue perfusion as the local transfer flux between the compartments. Thus enabling a clear distinction between fluid contributing to local perfusion and fluid in transit through the voxel.

Second, we formulate a hybrid model where a Darcy type porous media model is combined with two network models representing arterial and venous flow respectively. This model will be used in lieu of real observations to provide tracer-concentration data for the assimilation exercise as well as ground truth perfusion maps.

### 0.1 Porous media model used for parameter estimation

We model a region of live tissue as a three dimensional spatial domain Ω partitioned into four compartments. Three of the compartments constitute the vasculature, and are all represented as porous media flow models. The fourth compartment represent the extravascular tissue, and will for now be considered a passive tissue matrix confining our pore spaces.

Let the subscript *i* ∈{*a, v*} indicate the arterial or venous compartments respectively. As detailed in [19], we consider two porous flow models each governed by the Darcy constitutive relation 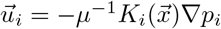, the incompressibility condition 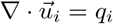, and conservation of contrast agent 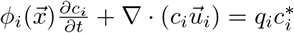. Here 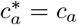, while in general 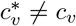, i.e. departing (*q <* 0) fluid carries the overall concentration of the compartment, while entering (*q >* 0) fluid in general carries a different concentration. The arterial and venous systems interact via a third porous medium, representing “small scale” vasculature. This compartment is given a simplified representation in terms of only a throughput conductivity 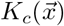 and a porosity 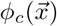, leading to fluxes

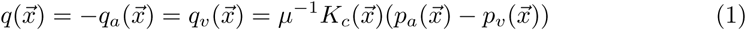

and transit times given by 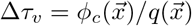, i.e., pore volume per throughput volume rate. While subscript *c* alludes to capillaries, the compartment will typically also capture other parts of the microvascular network like arterioles and venules. Although the capillary compartment per construction excludes any lateral transport, by regulating the throughput conductivity it may induce such transport in the arterial or venous compartments.

For model identification purposes, we consider eight parameter fields, three scalar porosity fields *ϕ*_*a*_, *ϕ*_*v*_ and *ϕ*_*c*_, two anisotropic diagonal permeability tensor fields *K*_*a*_ and *K*_*v*_, and one scalar permeability field *K*_*c*_. (For a 3D case there will be ten parameter fields.) Observation of time series for the contrast agent consentration on each voxel will guide the identification process. As outlined above, the local perfusion estimate for each voxel is taken to be the capillary flow rate *q* .

The spatial domain will be embedded in a regular Cartesian mesh, where selected cells will be made inactive to map out more complex geometrical structures. Boundary conditions will be a combination of pressure conditions and no-flow conditions, and the flow pattern will typically be driven by the difference between an arterial inlet pressure and a venous outlet pressure. Using a five (seven in 3D) point finite volume stencil, we solve for pressures *p*_*a*_ and *p*_*v*_, velocities 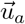 and 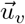, and transfer flux *q*.

To solve for the transport of contrast agent, we utilize that the formulation chosen for the fluid flow obeys a discrete maximum principle. Thus any two voxels connected by a streamline have a strict upstream-downstream relation, and the contrast agent advection can be computed sequentially voxel by voxel starting from the most upstream cell. We combine mass conservation 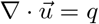 and time-of-flight *τ*, defined by 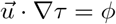 into 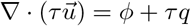, and consider a bounded control region of volume *V*. We partition the mass rate, *F*, through region boundaries according to flow entering (up) and leaving (dn):

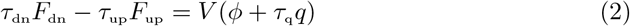

Thus one attribute an average arrival time, *τ*_up_ for fluid entering, and an average departure time, *τ*_dn_ for fluid leaving. Assuming the source term *q* to be uniform over the volume, the associated average time *τ*_q_ will either carry an upstream value (*q >* 0) into a venular compartment, or leave (*q <* 0) an arterial compartment with a value “somewhere”between *τ*_up_ and *τ*_dn_. For the computation in the current paper, we have lumped the transit times into a single quantity for each compartment, and write Δ*τ*_*v*_ = *V ϕ*_*v*_*/F*_dn,*v*_ for *q >* 0 (venular) and Δ*τ*_*a*_ = *V ϕ*_*a*_*/F*_up_,_*a*_ for *q <* 0 (arterial). This essentially means that we consider the flow to enter capillaries at the downstream end of an arterial compartment, and depart capillaries at the upstream end of the venular compartment.

The contrast agent concentration is sampled at discrete times of uniform increment Δ*t*, and the advection of the discrete contrast vector is carried out in three steps related to the arterial, capillary, and venous compartment respectively. Starting from the most upstream arterial voxel, the upstream concentration 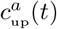 is shifted by transit time Δ*τ*_*a*_ to obtain the downstream concentration 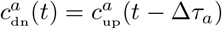. We also compute volume averaged concentration 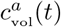, serving the dual purpose of arterial voxel concentration and input concentration for the capillary compartment. For multidimensional domains, the upstream concentration 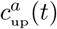 will generally be a flux weighted combination over several upstream neighbor connections, confer the lumping process leading to Eq 2. Similarly, by shifting and averaging using the capillary transit time Δ*τ*_*c*_, we obtain the capillary downstream concentration 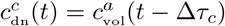 and the capillary volume average 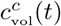. For the venular compartment, we again start from the most upstream voxel and for each voxel we compute a lumped upstream concentration vector 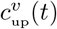 as a fluxweighted sum over upstream neighbor connections and the outflux 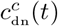 from the capillary compartment. Shifting by transit time Δ*τ*_*v*_ yields the downstream concentration vector 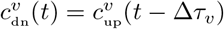. Finally the venous volume average concentration 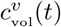 is computed, and combined into a total (including extravascular tissue) volume concentration for each voxel 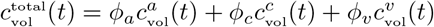.

### 0.2 A hybrid-scale flow model used to generate synthetic data

Here we consider an hybrid-scale model formulation, integrating one dimensional network flow with two- or three-dimensional continuum models. Known ground truth parameter values are assumed for this model when generating synthetic data. We presented the full details of this model in Hodneland et.al., [9], and here we will briefly recap some main points. Compared with the pure Darcy formulation presented in the preceding section, the arterial and venous compartments are now both augmented with flow networks representing visible arterial and venous vessel structure respectively, confer Fig 1.

**Fig 1.**
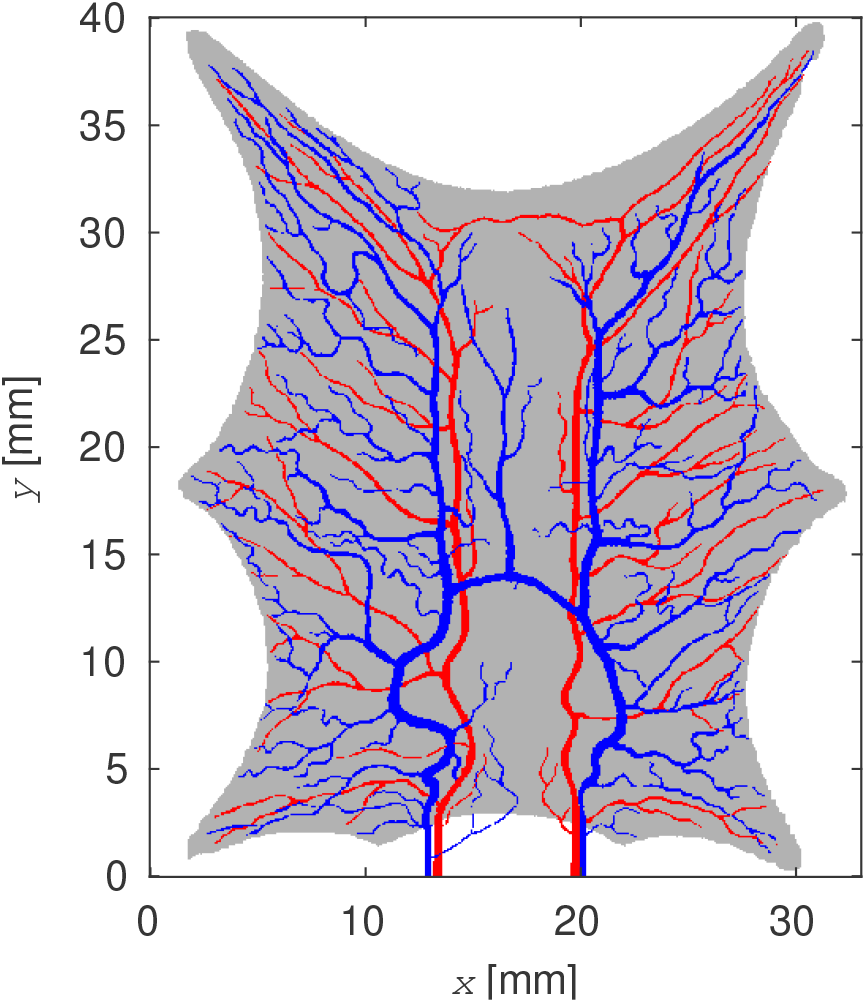
Vascular network for a stretched frog tongue. Arteries are red, and veins are blue. Gray color represent the tongue tissue. The inlets and outlets are at the lower edge.

The nodes of each network are classified as *interior, terminal*, and *root* nodes. A single node can have multiple roles. We associate a pressure value to each node location, and neighboring nodes are connected by edges representing vessel segments. For each edge we then have a one-dimensional flux that is related to node pressures via the Poiseuille law. We impose a mass balance condition at each node, requiring associated fluxes to satisfy net zero accumulation.

Each *root node* represents a cut-off for our region of interest, and we assign a pressure boundary conditions. Each *terminal node* represent a coupling to the associated continuum model, and we considered the flux to be proportional to the pressure difference between the node and the continuum at each node location. In order to provide for a smooth transition, a distribution functions in terms of a Gaussian kernel is centered at each location. This can be seen as accounting for missing structure in the transition between visible vessels represented by the the network, and the microstructure represented by the continuum, see [9] for details.

We combine the above with standard Darcy formulations for the arterial and venous continuums respectively, and introduce a coupling term between the two in terms of a flux proportional to local pressure difference. We identify this flux as the perfusion supplying the local tissue. We solve the total coupled formulation to obtain pressure and flux distribution across the combined model. Based on the fluxes obtained, we model advection of contrast agent. At each root node of the arterial network, we specify contrast concentration in the entering blood stream in terms of an arterial inflow function (AIF). At each *terminal node*, we again apply the Gaussian kernel to obtain a smooth distribution of entering contrast agent. Compared with the previous section where we used an explicit approach for agent transport, we here perform a fully implicit procedure based on a backward Euler discretization of the advection equation.

### 0.3 Bayesian inversion

The Bayesian method we use to assimilate tracer concentration data is introduced in [18]. This methodology has proven to be robust and efficient when assimilating big amounts of data to large-scale sub-surface reservoir models, see e.g [14, 15]. The technique is based on a regularized Levenberg-Marquardt method, and iteratively searches for the minimum of an average cost problem. The approach utilize an ensemble of model parameters, where each member of the ensemble is denoted *m*_*j*_, *j* = 1, ⋯, *N*, and *N* is the ensemble size. The unknown parameters are tissue properties such as transmissibility and porosity. Further, we denote the blood circulation model 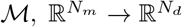 and the simulated concentration data are denoted *d*_*j*_ = ℳ(*m*_*j*_), *j* = 1, ⋯, *N*. The cost function of interest is defined as:

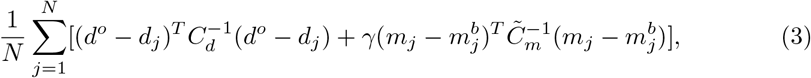

where *d*^*o*^ is the tracer concentration data and 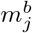 are given background (or best guess) parameters (defined below). The second term in the cost function is the regularization part, and *γ* is a (user defined) weight parameter. The data errors are assumed to be Gaussian distributed with zero mean and covariance *C*_*d*_. The matrix 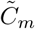 is the sample covariance of the background parameters.

The above problem is solved iteratively and, without going into details (see [18] for a full method description), we state the update formula for ensemble member *j* at iteration *i* + 1:

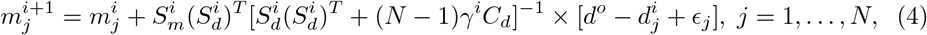

where 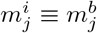 is adaptively updated. Initially these parameters are drawn from a user defined distribution (later referred to as the initial or prior ensemble). The columns in the matrices 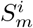 and 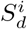 are given by 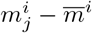 and 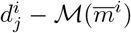, respectively, where 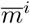 is the sample mean of the parameter vectors at iteration *i*. The perturbation terms *ϵ*_*j*_ are samples from the data error distribution. Similar to the background parameters, the weighting term *γ*^*i*^ is updated for each iteration. This parameter is increased if the average data mismatch increases, otherwise the parameter is reduced. In addition, a stopping criteria is selected such that the data mismatch of the final solution is of the same magnitude as the noise level.

#### 0.3.1 Localization

As mentioned in the introduction, inaccurate correlations between parameters and measurements makes it necessary to restrict the updates of parameters. We pursue a correlation-based localization technique [17], which is best described by formulating the update step (Eq 4) as

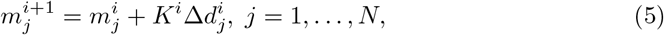

where

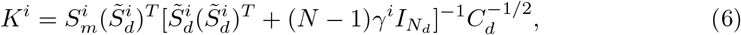

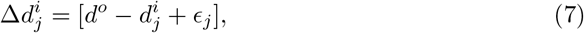

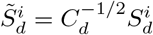 and 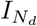 is the *N*_*d*_-dimensional identity matrix.

Correlation-based localization involves computation of a correlation matrix between observations and model parameters. In order to avoid huge memory requirements, we use a truncated singular value decomposition (TSVD) to project the data onto a subspace consisting of the dominant singular vectors. Applying TSVD to 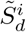 gives

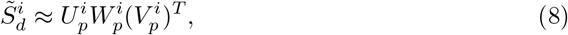

where *p < N* is the number of singular values after truncation. The matrix 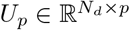 contains the left-singular vectors, *W*_*p*_ is a diagonal matrix composed of the singular values, and *V*_*p*_ ∈ℝ^*N ×p*^ contains the right-singular vectors. We keep 99.9 % of the sum of descending singular values. Substituting Eq 8 in Eq 5 gives

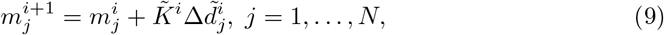

where 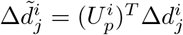 is the projection of the difference between simulated and real measurements, onto a subspace consisting of the dominant directions contained in *U*_*p*_. The matrix 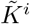 is the Kalman gain matrix after data projection. Introducing a tapering matrix Λ^*i*^, we define the update step

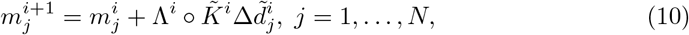

where ∘ denotes the matrix Schur product. The tapering matrix is either 1 or 0, and is (at a given iteration *i*) computed based on information about correlations between the projected data innovations 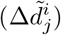 and the model parameters 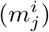. The details for computing the tapering matrix is not included here, and we refer to [17] for a full description. Examples of localization areas are given below.

## Results

In the current work we illustrate the methodology on a two dimensional problem. The example is based on an image of a frog tongue found in the book by Cohnheim [4], see Figure 1. The frog tongue is stretched and pinned to the surface at six points, so that the thickness is sufficiently small for visual identification of major arteries and veins. The dimension of the domain we have set to be 33 mm in the x-direction, and 40 mm in the y-direction. This data has the advantage of being truly 2D but still biologically relevant. Selecting as 2D slice from a 3D geometry will not manage to account for out-of-plane fluxes. The measurements are synthetically generated using the hybrid-scale model, and the unknown parameters are estimated using the porous media model. The code and data are available at [16].

With frequent sampling of data in time and space, the amount of data would be excessive. Therefore, only a fraction of the data are used. Since the data will be correlated in time, an algorithm for finding a reasonable reduced set of data is designed. The algorithm is designed to use a fraction of the points in time (i.e., it uses whole images for the selected time points). In the case study described below, we decided to use one-tenth of the available time points. In our case the hybrid-scale model is sampling data every second for a time interval of 150 seconds. To select the time points we form one vector *x*_*i*_ for each time point which we consider as a stochastic variable. Then we form the covariance matrix *C* = Cov(*x*_*i*_, *x*_*j*_) where 1 ≤*i, j* ≤*n*_*t*_ and *n*_*t*_ = 150 denotes the number of time points where data are sampled. Let *I* be a set of indexes from the set {1, ⋯, *n*_*t*_} and let *C*_*I,I*_ denote the principal submatrix of *C* containing the rows and columns from the set *I*. Motivated by D-optimal design [20] we would like to select *n*_*p*_ = *n*_*t*_*/*10 points such that det *C*_*I,I*_ is maximized over all sets *I* having *n*_*p*_ distinct entries from the set {1, ⋯, *n*_*t*_}. (For clarification: ⌊· ⌋ denotes the floor operation, 10 is our choice, and might obviously be changed.)

When *n*_*t*_ is increasing, the maximization problem described above would be very time consuming to solve by a brute force search. Since solving this optimization is only done to improve the speed and efficiency of the Bayesian inversion, we use have designed a greedy approach to find a reasonable solution to the problem of finding the set *I* which maximize det *C*_*I,I*_ over all sets *I* of size *n*_*p*_. The greedy algorithm is designed by first selecting *k*_1_ such that 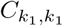 is maximized. This will be the solution to the maximization problem if the size of the set *I* is 1. From this we start to form sets *I*′ of increasing size until we reach the size *n*_*p*_ by adding one index in each step. In the second step we find *k*_2_ with *k*_1_ fixed from the first step such that det *C*_*I*′_, _*I*′_ where *I*′ has size 2. For an arbitrary step we are given indexes *k*_1_, ⋯, *k*_*j*−1_ and find *k*_*j*_ maximizing det *C*′_*I*_, _*I*′_ where *I* ′= {*k*_1_, ⋯, *k*_*j*−1_} ∪ *k* where *k* is any of the indexes 1 ≤ *k* ≤ *n*_*t*_ different from the previous selected indexes *k*_1_, ⋯, *k*_*j*−1_. The algorithm terminates when the size of *I*′ is *n*_*p*_.

The experimental setup (“truth”) for the hybrid-scale model is given in Table 1. The model is run on a grid with dimension 515 *×* 634, and data are contrast agent concentration values in every gridcell. As a base case, the simulated measurements are upscaled using a stencil of 4 *×* 4 grid-cells. In order to limit the amount of data we down-sample the measurements as described above. This gives a total of 191220 values. The blood stream entering the domain (AIF) is represented by a Gamma-variate function, see Fig 2.

**Table 1.** Experimental setup for the hybrid-scale model. See [9] for more information about the model.

| Parameter | Symbol | Unit | Value |
| --- | --- | --- | --- |
| Arterial porosity | $\phi_a$ | - | 0.05 |
| Venous porosity | $\phi_v$ | - | 0.1 |
| Arterial permeability | $K_a$ | m<sup>2</sup> | $1 \cdot 10^{-13}$ |
| Venous permeability | $K_v$ | m<sup>2</sup> | $5 \cdot 10^{-13}$ |
| Inlet pressure | $p_a^0$ | kPa | 10.6 |
| Outlet pressure | $p_v^0$ | kPa | 1.60 |

**Fig 2.**
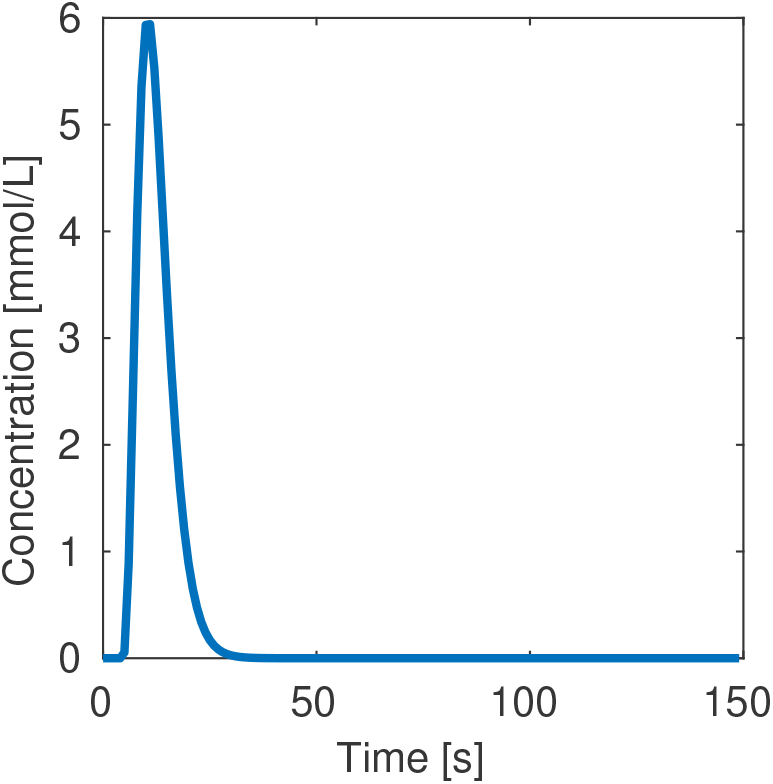
Arterial input function.

In this example study we select to estimate transmissibilities and porosities in the arterial, venous, and capillary compartments. In the arterial and venous compartments we estimate transmissibility in both the x- and y-direction. This gives a total of eight unknown parameter fields. Initial values are generated as Gaussian random fields with given mean and Gaussian variograms. We assume constant mean values in the frog tissue. In the arteries and veins we use transmissibility (in both x- and y-direction) given by *T* = *D*^2^*/*32*μ*, where *D* is the vessel diameter. This relation is found by comparing the Darcy equation (with porosity equal to 1) with the Hagen-Poiseuille equation for laminar incompressible flow. The variogram ranges are drawn from a Gaussian distributions with mean values equal to 74 and 60 grid-blocks in the x- and y-direction, respectively. The variance (in both directions) is equal to one grid-block. The relatively long variogram ranges are based on an assumption that the tissue has an approximately homogenous structure. To ensure that all parameters are physically reasonable we also impose upper and lower bounds for the values. All transmissibility fields are transformed using the natural logarithm. The statistical parameters are listed in Table 2. The porous media flow model is discretized using 128 cells in the x-direction, and 158 cells in the y-direction (base case). There are 7507 inactive cells that fall outside the boundary of the frog tongue. Each parameter field then consist of 12717 values, and the total number of parameters is *N*_*m*_ = 101736. We have also listed the true parameters used to generate the measurements with the hybrid-scale model. The viscosity, boundary pressures and AIF function are identical for the hybrid-scale model and the porous media model.

**Table 2.** Statistical parameters used to generate the initial ensemble. Lower and upper bounds are denoted lb and ub, respectively. Note that the mean transmissibility values are only valid in the frog tissue (outside the identified main vessels). Inside the vessels, the transmissibilities are given by the Hagen-Poiseuille equation.

| | $\mu$ | $\sigma$ | lb | ub | true |
| --- | --- | --- | --- | --- | --- |
| $\phi_a$ | 0.1 | 0.1 | 0.001 | 0.999 | 0.05 |
| $\phi_v$ | 0.1 | 0.1 | 0.001 | 0.999 | 0.1 |
| $\phi_q$ | $10^{-4}$ | $10^{-5}$ | $10^{-6}$ | 0.1 | NA |
| $\ln T_a^{x,y}$ | -20.7 | 1 | -26 | -10 | -20.7 |
| $\ln T_v^{x,y}$ | -20.7 | 1 | -26 | -10 | -19 |
| $\ln T_q$ | -16.8 | 1 | -26 | -8 | -13.4 |

On Fig 3 we show three realizations from the prior distribution for y-transmissibility in the arterial compartment, and the porosity in the venous compartment. Note that the blood vessels are not visible on the porosity fields, because the mean porosity values are equal (0.1) both inside and outside the vessels. This value is adopted because it is used in the hybrid-scale model when generating the measurements, and the motivation is to avoid porosity values larger than 1 in the parts of the tissue where arteries and veins are crossing (on top of each other). Examples of the computed localization domains based on the initial ensemble, 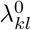, for selected observations, are shown on Fig 4. The areas change for each projected observation (*l*), and at every iteration (*i*). The initial ensemble is used as starting values for the algorithm given by Eq 4. The maximum number of iterations is set to 10, and in addition the algorithm will stop if the relative change of average data mismatch is less than 10%. In this example five iterations were always performed before the method terminated.

**Fig 3.**
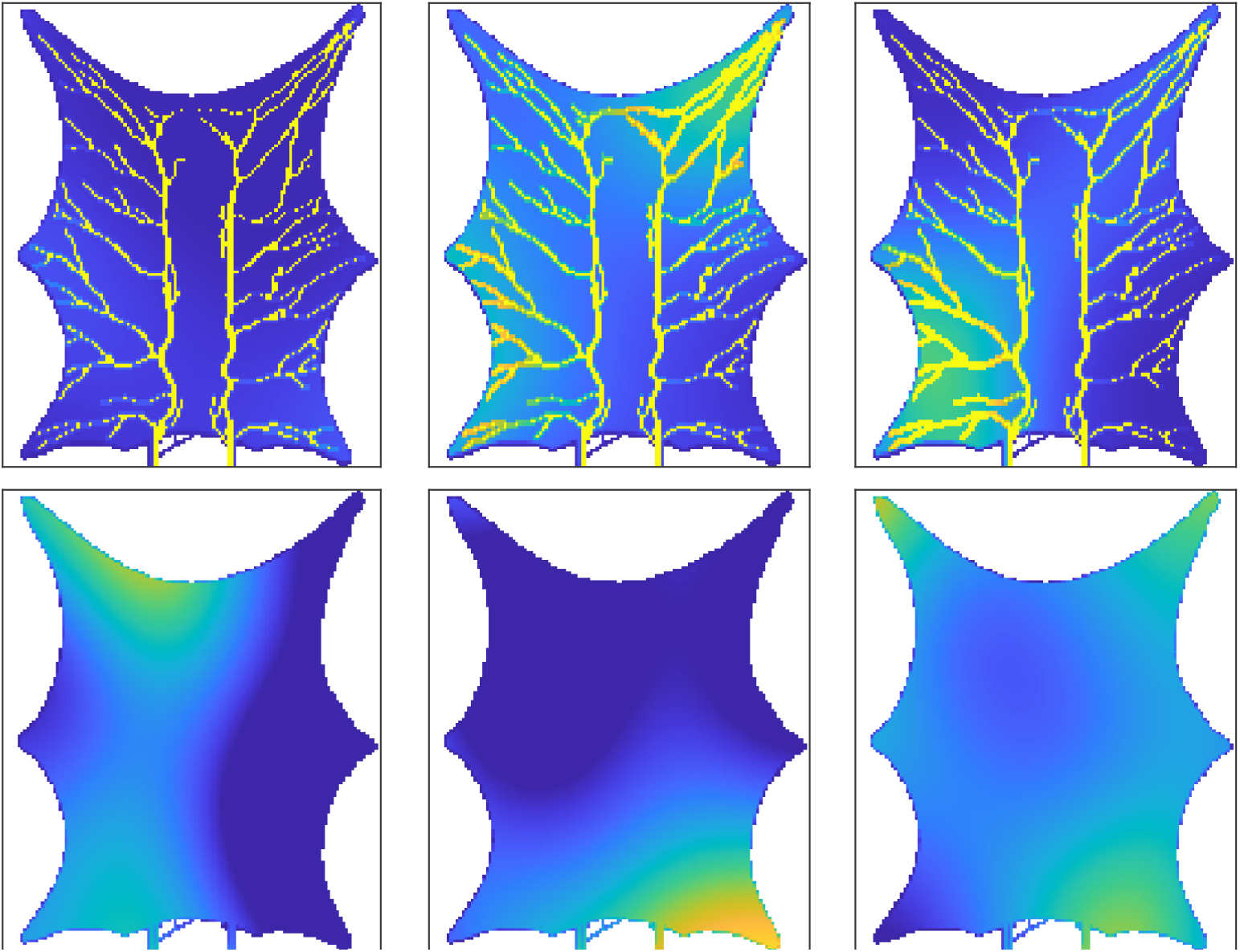
Three initial ensemble members for y-transmissibility (m^2^) in the arterial compartment (*top*) and porosity (m^3^/m^3^) in the venous compartment (*bottom*).

**Fig 4.**
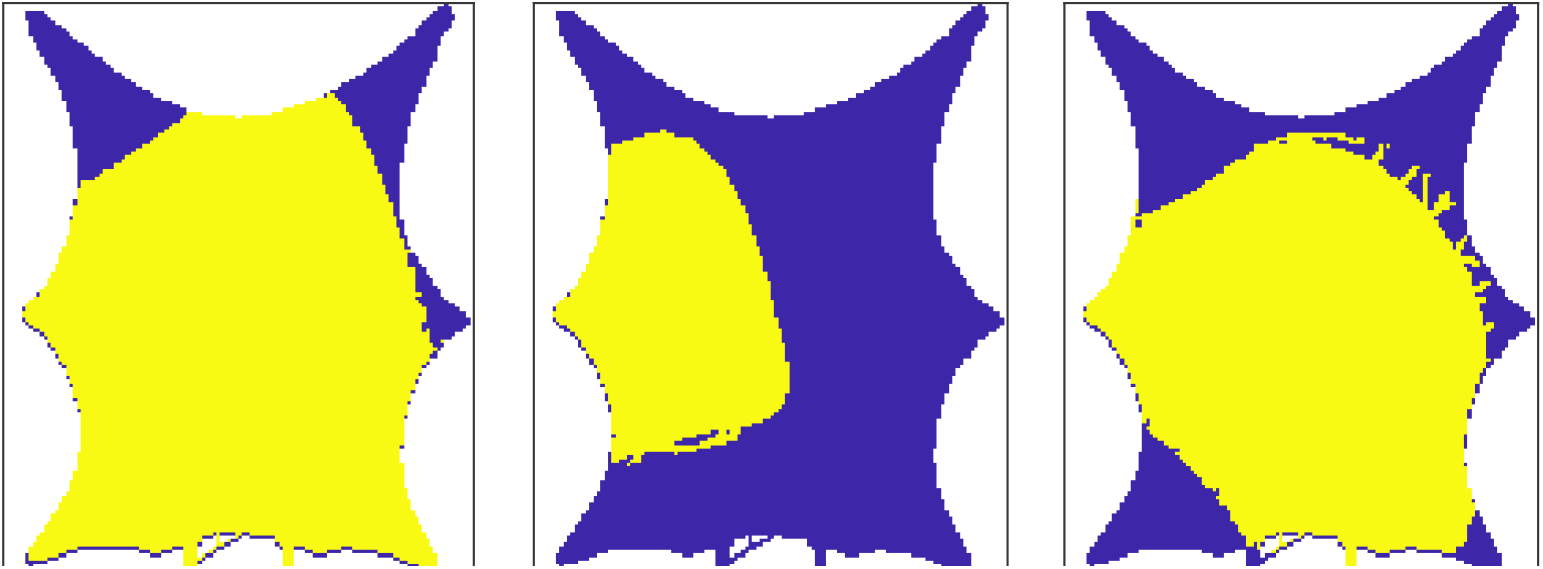
Correlation-based localization area for three observations (*l* = 1, 3, 8), using the initial ensemble (*i* = 0). The yellow area represent positions where parameters are updated (Λ^0^ = 1).

### 0.4 Estimated perfusion

The estimated perfusion follows from Eq 1. The true perfusion (see [9], Equation 7) is shown on the left picture on Fig 5. The perfusion range is between 5 mL/min/100mL and 21 mL/min/100mL. The spatial relative error in estimated perfusion is shown on the middle and right pictures. The relative error is computed as (*P*_*T*_ − *P*_*E*_)*/P*_*T*_, where *P*_*T*_ is the true perfusion and *P*_*E*_ is the estimated perfusion. The prior and posterior errors are calculated by simulating the perfusion using the mean of the prior and posterior ensembles. Note that the perfusion is zero in the vessels, and the vessels are colored for enhanced visibility. There is a clear improvement for the posterior estimate, and the relative error is especially reduced in the lower part of the domain, where the prior values are as low as −14. In particular, large errors occur when the true perfusion is low. The relative error using the posterior estimate is minimum −2.6. From the results we see that both the prior and posterior overestimate the perfusion in most of the domain. In order to further quantify the accuracy of the estimated perfusion we compare the results with two widely used techniques within radiology: the maximum slope method and deconvolution based on singular value decomposition. We will not present these methods here, but refer to [6] for readers interested in details. On Fig 6 (left) we show the average perfusion (*P*) computed for the entire frog tongue domain. The true value is shown in blue and is 9.5 mL/min/100mL. The maximum slope method and deconvolution method slightly overestimates the perfusion value (the error is 1.1 mL/min/100mL), whereas the prior model returns a value as high as 71 mL/min/100mL. The posterior model obtained after assimilation of concentration data is 8.9 mL/min/100mL, slightly below the true value. On the right plot we show corresponding results for four regions, obtained by dividing the domain into equally sized parts. The posterior model performs best in all regions. The estimated perfusion results are generalized on Fig 7, where the number of regions are successively increased. More accurately, we compute perfusion on the following grids: 1 *×* 1, 2 *×* 2, 4 *×* 5, 8 *×* 10, 16 *×* 20, 32 *×* 39, 64 *×* 79, 128 *×* 158. The most refined grid is the base case. For each sub-division of the domain we compute the perfusion for all regions and calculate the mean values for the errors. I.e., for a specific division of the domain into *s* regions of interest we compute the mean absolute error (MAE) as

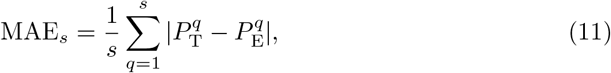

where 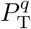 is the true perfusion in region *q* and 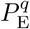 is the estimated perfusion in region *q*, using one of the estimation (E) techniques: maximum slope, deconvolution, prior model, or posterior model. In the above calculations we utilize the information about the position for arteries and veins by setting the perfusion value in a region to zero if the region contain more than 50 % vessels. The maximum number of regions are all voxels in the base case domain (128 *×* 158), excluding the voxels that contain arteries or veins. The conclusion from this figure is that the posterior model is better than all the other techniques we have considered, for all partition levels.

**Fig 5.**
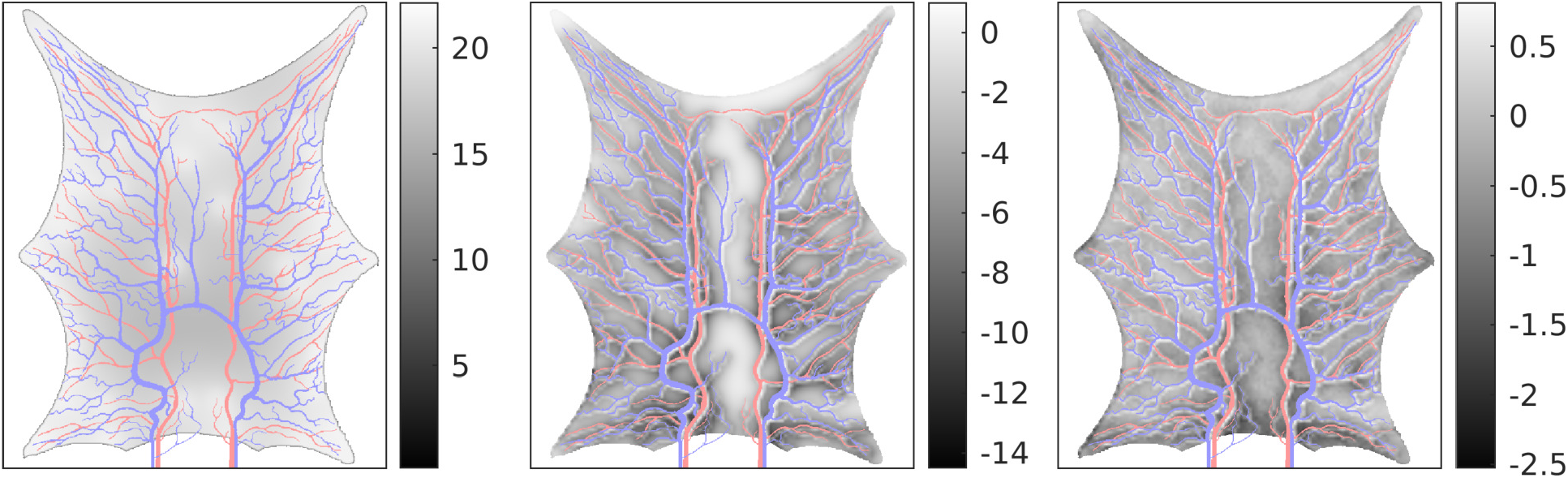
*Left:* True perfusion (mL/min/100mL). *Middle:* Relative error in simulated perfusion using the mean prior ensemble. *Right:* Relative error in simulated perfusion using the mean posterior ensemble. Note the different ranges on the relative error maps.

**Fig 6.**
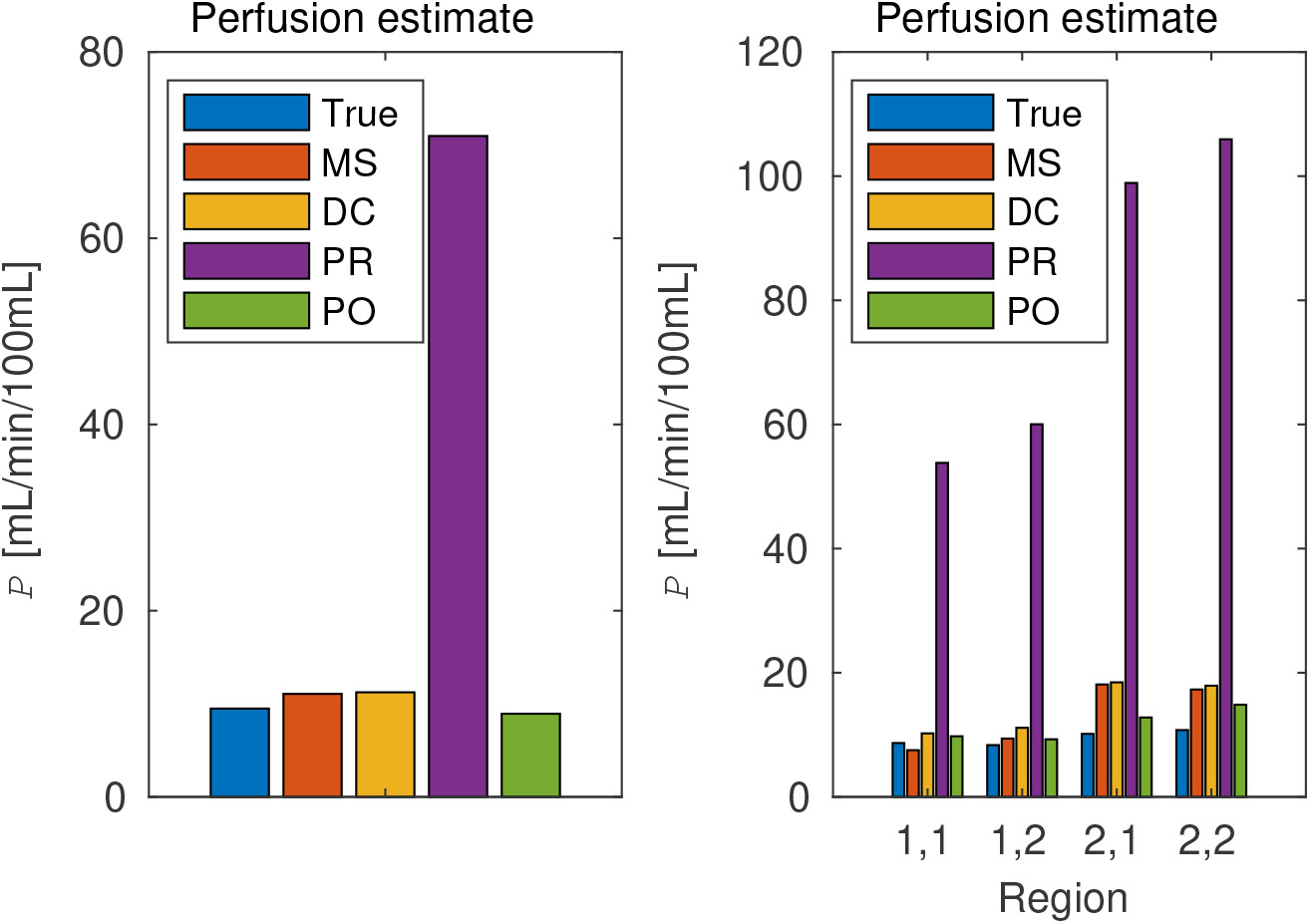
Comparison of estimated perfusion using different techniques. On the bar plots, ‘MS’ is the maximum slope results, ‘DC’ is deconvolution results, ‘PR’ is results using the prior model, and ‘PO’ are results using the posterior model after data assimilation. *Left:* entire domain. *Right:* four sub-regions. On the x-axis, *i, j* denote the region and *I* is the index in the x-direction (from left to right) and *j* is the index in the y-direction (from top to bottom).

**Fig 7.**
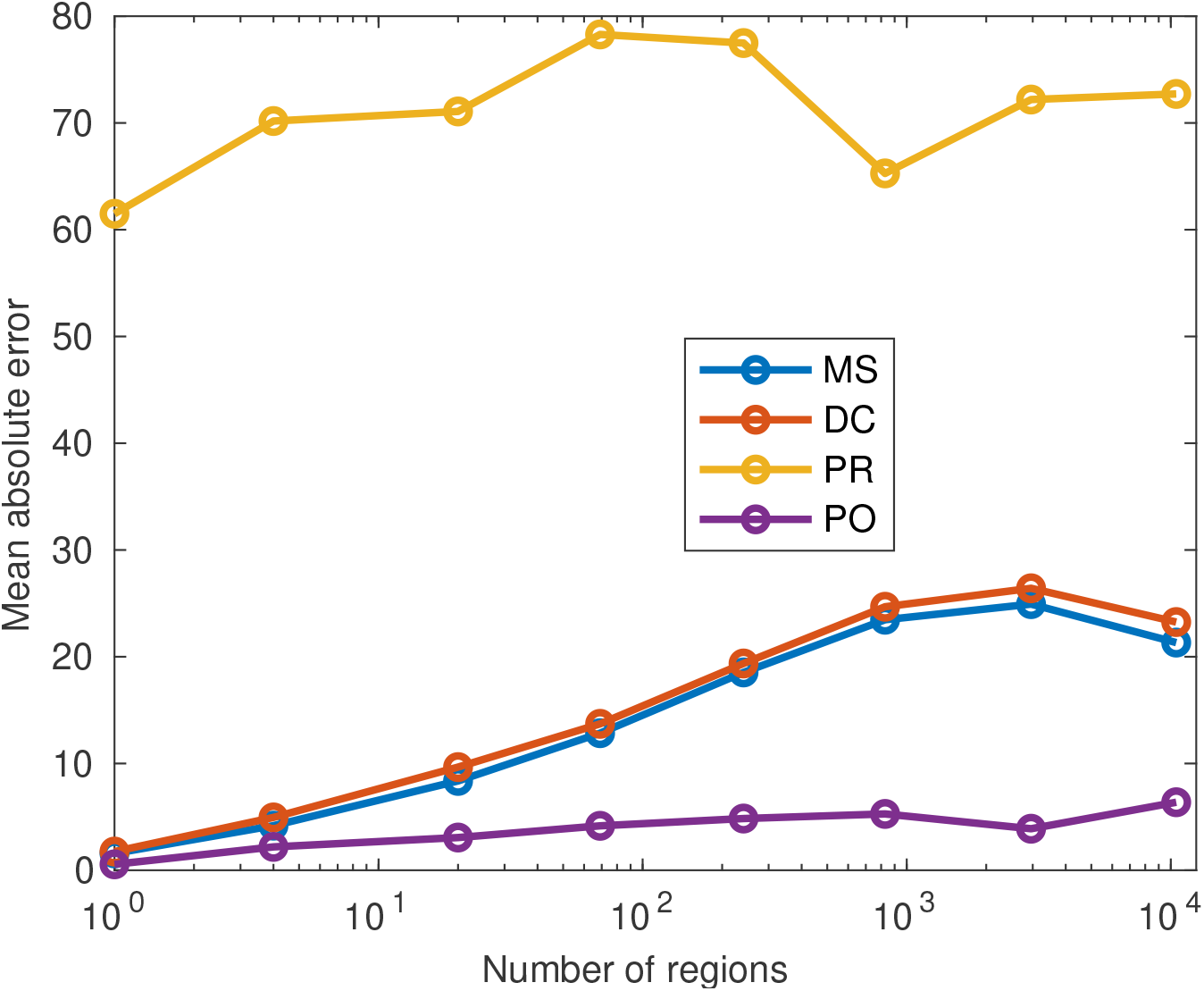
Mean absolute error for estimated perfusion for increasing number of regions. On the plot, ‘MS’ is the maximum slope results, ‘DC’ is deconvolution results, ‘PR’ is results using the prior model, and ‘PO’ are results using the posterior model after data assimilation. The x-axis is logarithmic, and the range is from 1 to 10483.

### 0.5 Estimated concentration

Fig 8 shows the spatial distribution of contrast concentration at four points in time, simulated using the full resolution with 128 *×* 158 gridcells. At time equal to 33 seconds a clear concentration pattern is seen in the data. The prior estimate illustrate the fact that the blood vessels in the porous media model is not sealed, and there is a diffusion through the vessel walls. This phenomenon is reduced in the posterior model, but the the circular concentration areas seen in the data are not recovered. The rightmost figure shows the standard deviation for the posterior ensemble. As explained in the introduction, it is preferable to avoid underestimation of the uncertainty after iterating. The standard deviations are higher than zero, especially early in the time period, thereby indicating that collapse of the ensemble is avoided. Visually, we obtain improvements for the posterior concentration at all points in time, compared to the prior concentration values. The estimated concentration is further visualized at Fig 10. This figure shows concentration curves at nine positions (indicated on Fig 9) in the frog tongue. These positions are evenly distributed in the domain, and among the points, the middle-right is in a high permeability zone (blood vessels). Clear reduction in the uncertainty is seen for the posterior curves (shown in green), compared to the prior curves (shown in blue). The posterior curves are of varying quality with clear improvements at e.g. at the lower-right voxel, but less accuracy at e.g. the middle voxel. Despite inaccuracies at some points in time, the overall data mismatch is reduced from approximately 2.8 · 10^5^ to approximately 9.0 · 10^4^, as seen on Fig 11. The values are shown on a logarithmic y-axis for better visibility. The formula used to compute the data mismatch for ensemble member *j*, at iteration *i*, is

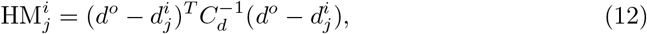

where the measurements (*d*^*o*^) and simulated observations 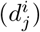 are concentration data.

**Fig 8.**
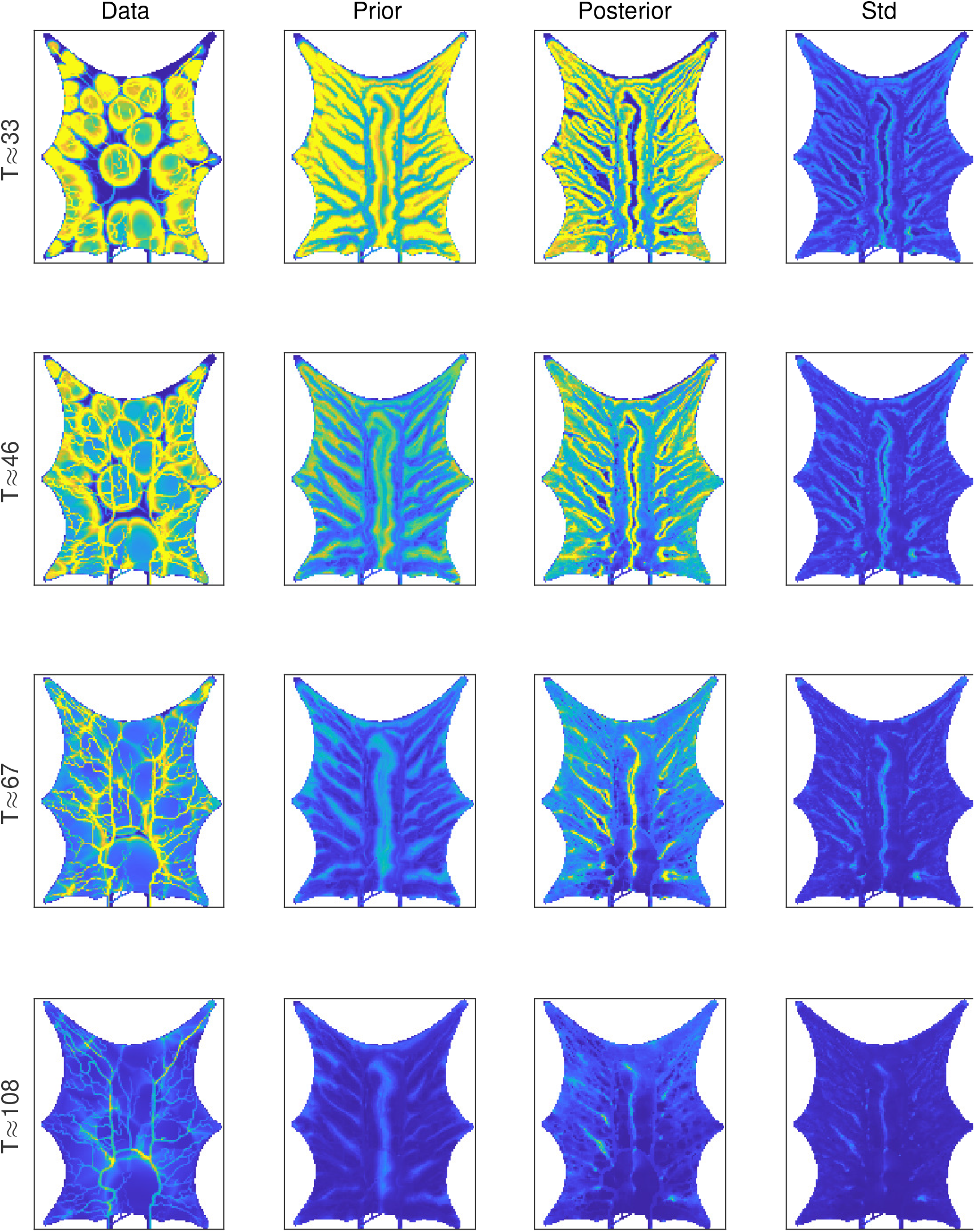
Spatial distribution of contrast concentration at four points in time. Blue color indicates low concentration and yellow indicates high concentration. The same scale (colormap) is used for all plots. The columns shows from left to right the data, the prior mean, the posterior mean and the posterior standard deviation.

**Fig 9.**
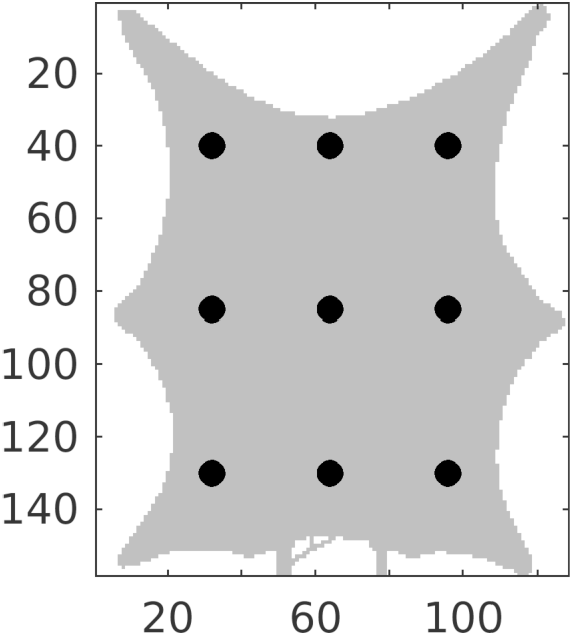
Positions (shown as black dots) used to visualize concentration curves. The grid indices are shown on the axes.

**Fig 10.**
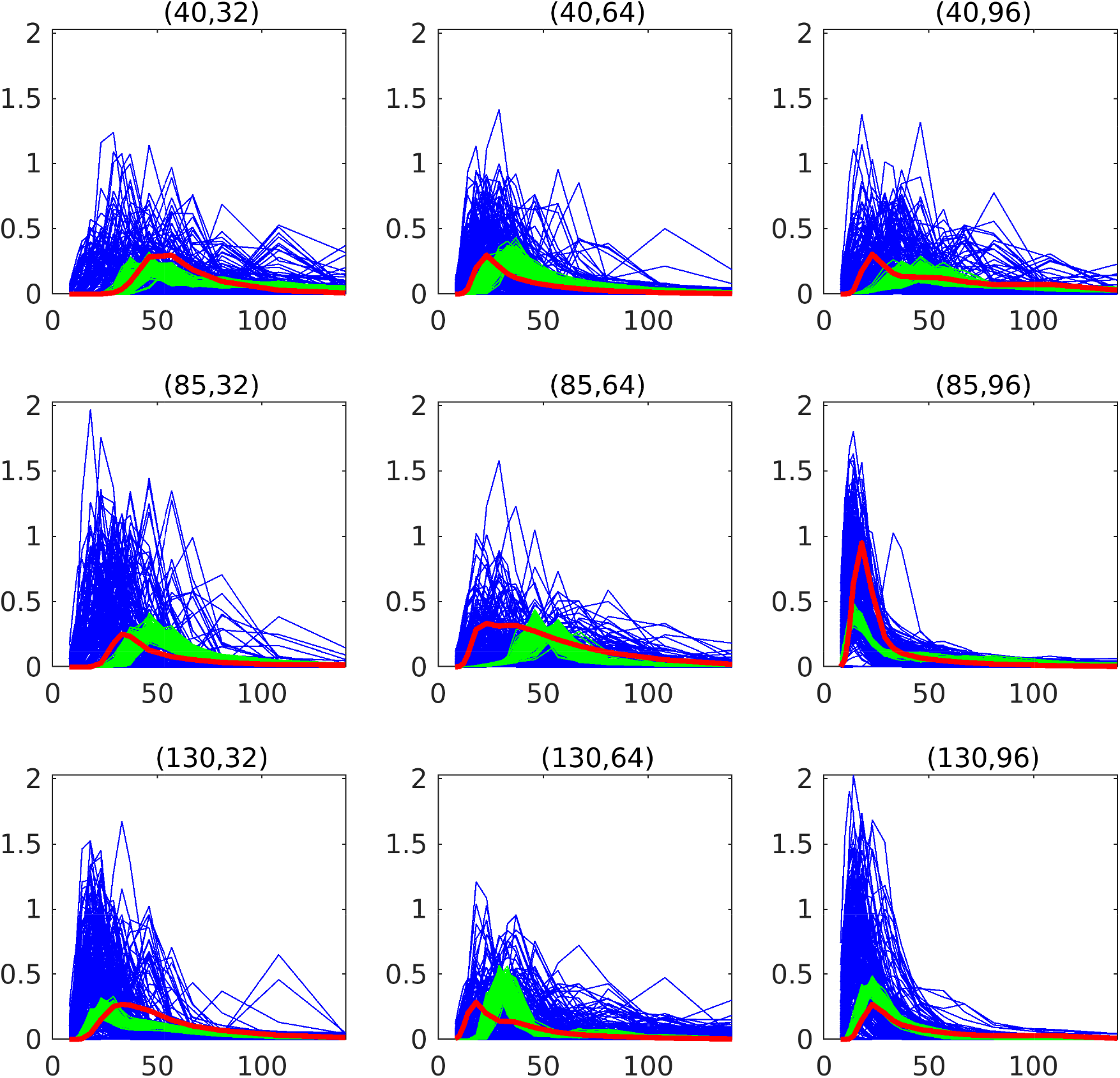
Temporal distribution of contrast concentrations at nine positions, given by the (coarse) grid indices in the titles. Red color indicates the measurements, blue color represent the initial prior ensemble, and green color represent the updated posterior ensemble. The x-axis is the time in seconds, and the y-axis is the concentration values in mmol/L.

**Fig 11.**
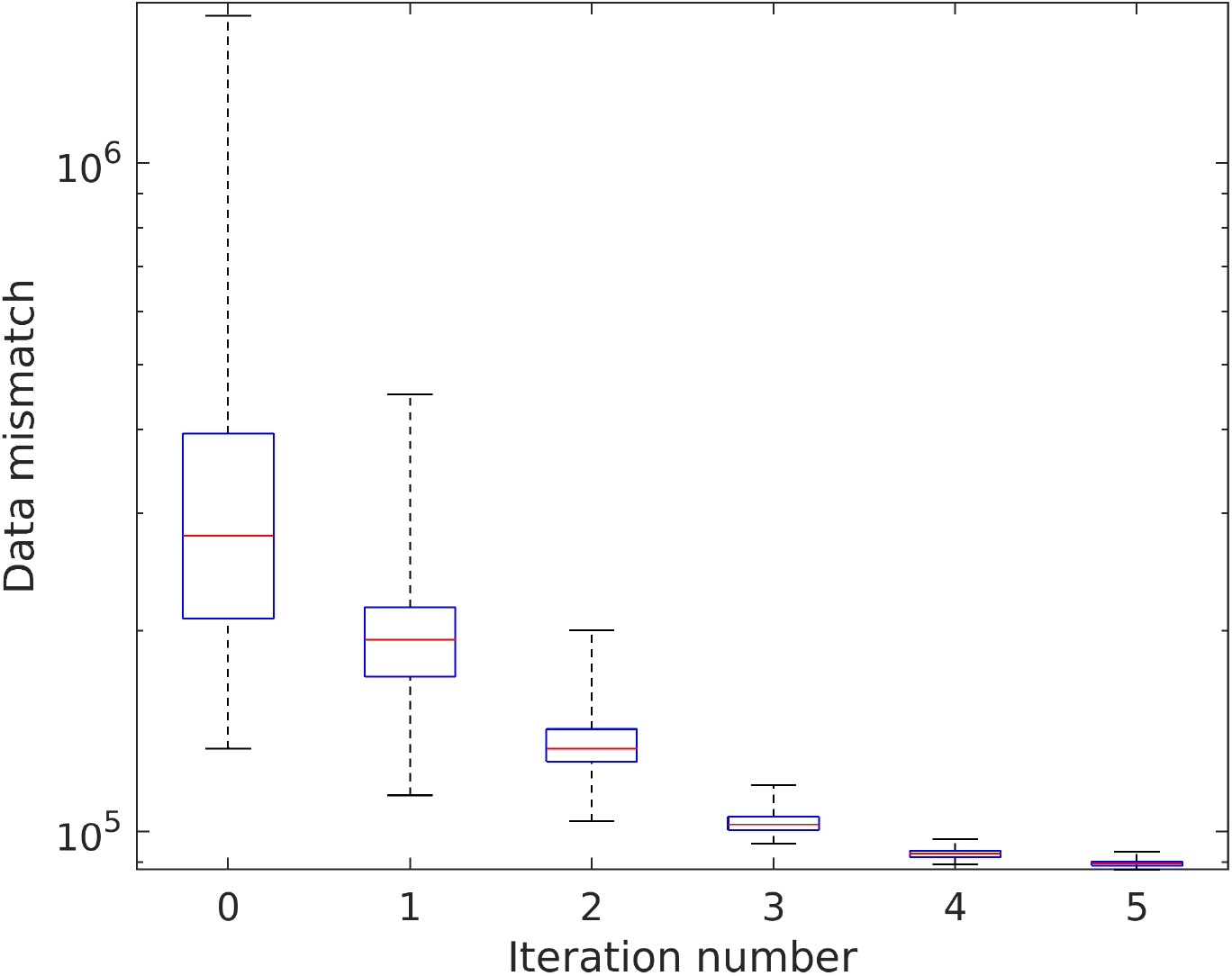
Data mismatch for concentration data. The y-axis is logarithmic. The red horizontal lines indicate the medians, and the blue horizontal lines indicate the 25th and 75th percentiles. The whiskers cover the extreme values.

## Discussion

We have demonstrated use of ensemble-based estimation techniques for assimilating processed contrast agent concentration data from dynamic MRI acquisitions into models for blood perfusion in organs. The methodology is applied to a domain with known vascular structure, and the realism of the methodology is enhanced by selecting different models for synthetic data generation and data assimilation. The data-generating hybrid-scale model includes a detailed representation of both the arterial and venous vessel structure, and a distribution function is used for the flux from the vessel network to the continuum. The porous media model used for data assimilation is simpler, and needs to represent vessel structures as high-permeable channels. We show that improved perfusion estimates are obtained, compared to traditional methods (maximum slope and deconvolution methods). The major contribution of our work is the ability to provide better estimates of the perfusion in any region of interest embedded in the organ of investigation. The importance of accurate estimates of perfusion is, as mentioned in the introduction, wide-ranging.

The work presented here represent a first step towards a full-scale tool for clinical usage. The next step is to evaluate the methodology on synthetic three-dimensional contrast data, and eventually real MRI-based measurements. When using real measurements there are challenges related to how the concentration data are scaled and pre-processed, that must be addressed. It has recently been suggested that adding in-silico trials would be helpful for development of better treatment of acute ischemic stroke [11–13] and new modeling approaches to handling different issues are being developed [5]. Obviously, patient specific models that align with observations would increase the value of in-silico models even further.

## Acknowledgments

This work is supported by the Norwegian Research Council project 262203 “Flow-based interpretation of Dynamical Contrast Enhanced Imaging data”.

## Notes

### Competing Interest Statement

The authors have declared no competing interest.

